# Restoration of high-sensitivity patterned vision in motion with an engineered light-gated G protein-coupled receptor

**DOI:** 10.1101/2022.04.07.487476

**Authors:** Amy Holt, Michael H. Berry, Jamie Lo, Prashant Donthamsetti, Meike Visel, Johannes Broichhagen, John G. Flannery, Ehud Y. Isacoff

## Abstract

Inherited retinal degenerations (IRDs) result in blindness due to apoptotic cell death of rods and cones, but spare other retinal neurons, providing a potential that delivery of a light-activated signaling protein to surviving neurons may restore vision. We previously demonstrated that aspects of vision could be restored by introduction into surviving cells of a G protein-coupled receptor for glutamate (mGluR) bearing a tethered photoswitchable agonist. However, this system, containing one photoswitchable agonist per glutamate binding site, yielded low sensitivity, responding only to visual stimuli at the intensity of bright outdoor light, similar to channelrhodopsins. To increase sensitivity, we designed a multi-branched photoswitch, bearing four light-activatable glutamates for each glutamate binding site. When tethered to a modified mGluR2 expressed in retinal ganglion cells *via* intravitreal AAV gene delivery, this photoswitch boosted sensitivity by ~100-fold compared to the unbranched (single photo-ligand) photoswitch. This improvement in sensitivity enabled an IRD mouse model (*rd1*) to perform visually-guided object recognition under incidental room light and pattern recognition using standard LCD computer displays. The restored line pattern differentiation approached the acuity reported for normal mouse vision. Pattern recognition functioned as well as wildtype vision with line patterns moving at speeds of up to 36°/s. In summary, this two-component chemical-optogenetic approach combines high sensitivity and high acuity with superior motion vision, and, unlike optogenetic gene therapy, can be adjusted for dose, upgraded, as new photoswitches are developed, and discontinued at will.

## Introduction

Blindness from most inherited retinal degenerations (IRDs) results from degeneration of rod and cone photoreceptors, but spares other retinal cells, leading to the opportunity that expression of light-activated signaling proteins in the surviving retinal neurons could restore vision. Light-activated ion channels, including microbial channelrhodopsins^1–8^ and the chemically engineered ionotropic glutamate receptor LiGluR^9,10^ have been used to restore aspects of vision to animal models of RP, and the channelrhodopsin ChrimsonR was recently shown to restore object detection in a blind patient^11^. These light-activated channels have fast kinetics but low sensitivity. Recent improvements in channelrhodopsin sensitivity have enabled vision restoration at moderate light levels in mouse and rat IRD models^12,13^, but ChrimsonR, which is also more sensitive than wildtype ChR2, requires intensifying goggles to provide useful vision in a patient^11^. One alternative approach is to repurpose native light-activated G-protein coupled receptors (GPCRs) in the retina, as these may operate at lower expression or luminance levels because they activate channels downstream of an amplifying signal cascade. The native opsins of rod photoreceptor cells-rhodopsin and those of the intrinsically photosensitive retinal ganglion cells-melanopsin, when ectopically expressed in retinal ganglion cells can restore a light response with 2-3 order of magnitude greater sensitivity than channelrhodopsins. These opsins enable animal models to distinguish light from dark under very dim lighting conditions^4,14–18^. However, when expressed in retinal neurons other than photoreceptor cells, they deactivate slowly and recover slowly. The faster of the two, rhodopsin, does not support patterned vision, even with an immobile visual stimulus, likely due to its slow kinetics^19^. In contrast, medium wave cone opsin expressed in RGCs provides the same sensitivity as rhodopsin, but with faster kinetics resulting from a more rapid shutoff rate, and does support line pattern recognition, not only with immobile images but also in slowly moving images^19^.

In an effort to combine high sensitivity with fast kinetics and restore visual responses to rapidly moving, patterned stimuli at indoor light intensities, we developed a 2-component strategy that combines expression in the retina of an N-terminal engineered glutamate-activated GPCR, fused to a self-labelling SNAP-tag, SNAP-mGluR2, which is sensitized to light by a synthetic photoswitch, BGAG_12,460_^10^. In previous work, a single BGAG_12,460_ attached to SNAP-mGluR2 expressed in retinal ganglion cells (RGCs) restored vision, but had inadequate light sensitivity, requiring line patterns to be displayed at high intensity to be detected^10^.

To increase light sensitivity, we use a redesigned BGAG_12,460_ with four times the number of light-activated azobenzene-glutamate ligands around each glutamate binding site of the receptor. When conjugated to SNAP-mGluR2 in HEK293 cells co-expressing the G protein gated inward rectifier (GIRK) potassium channel, this 4-branched version (^4x^BGAG_12,460_) approximately doubles the efficacy of GIRK activation compared to the unbranched (single photo-ligand) BGAG_12,460_^20^. Here, we find that branched ^4x^BGAG_12,460_:SNAP-mGluR2 in the RGCs of the highly visually impaired *rd1* mouse generates a light response with far greater sensitivity than single armed BGAG_12,460_. Formulation of ^4x^BGAG_12,460_ in β-cyclodextrin extends the restoration of light perception for 6 weeks after a single intravitreal injection. Following expression of the ^4x^BGAG_12,460_:SNAP-mGluR2 in the RGCs, the visually impaired *rd1* mice can now perform novel object recognition under incidental room light and distinguish between visual patterns displayed on a standard LCD computer screen. The acuity level of the restored line pattern differentiation approaches the theoretical limit for wildtype mouse vision. Moreover, line pattern recognition works as well as wildtype vision when patterns on visual displays move at up to 36o/s.

While the requirement of repeated administration of the BGAG chemical photoswitch adds an additional step to the gene therapy, it also provides the unique possibilities to adjust the dose, to upgrade as new photoswitches are developed and to be discontinued in case of undesired experience for the patient.

## Results

### Branched BGAG restores fast light responses to the *rd1* retina

We expressed a modified mGluR2 in RGCs that has which a SNAP-tag is fused to the N-terminal of mGluR2 (SNAP-mGluR2)^10^. Our synthetic photoswitch, **<u>b</u>**enzyl**<u>g</u>**uanine-**<u>a</u>**zobenzene-**<u>g</u>**lutamate (BGAG), attaches to the SNAP via a selective, bioorthogonal reaction of its *O*^6^-benzylguanine (BG) end by means of a covalent bond. The BG is connected to a photoisomerizable azobenzene-glutamate moiety (AG) via a linker made of n = 0-28 polyethylene glycol (PEG) repeats (BGAG_n_). BGAG with a classical (unmodified) azobenzene^21^ photoisomerizes in near-UV light (380 nm) from its dark-stable *trans* state, in which the glutamate is obstructed (unable to bind to the receptor’s ligand binding pocket) and the receptor is inactive, to its *cis* state, in which the glutamate is exposed, binds and activates the receptor. This BGAG returns to the *trans* state very slowly (hours) in the dark but can be rapidly (ms) photoisomerized to *trans* by blue-green light (488-532 nm) (BGAG_n,380/532_). A more electron-rich azobenzene photoisomerizes to *cis* under blue light (460 nm) and returns rapidly to *trans* in the dark^22^. For vision restoration, we therefore turned to BGAG_460_. We used a 12 PEG linker (BGAG_12,460_) since this linker length provides maximal photo-activation of SNAP-mGluR2^21^. We turned from a single armed BGAG_12,460_, in which each mGluR2 subunit of the dimer has one SNAP, and therefore one azobenzene-glutamate (AG), to a four-branched, ^4x^BGAG_12,460_, which bears four light-activated glutamates per subunit (**Figure 1A**).

**Figure 1:**
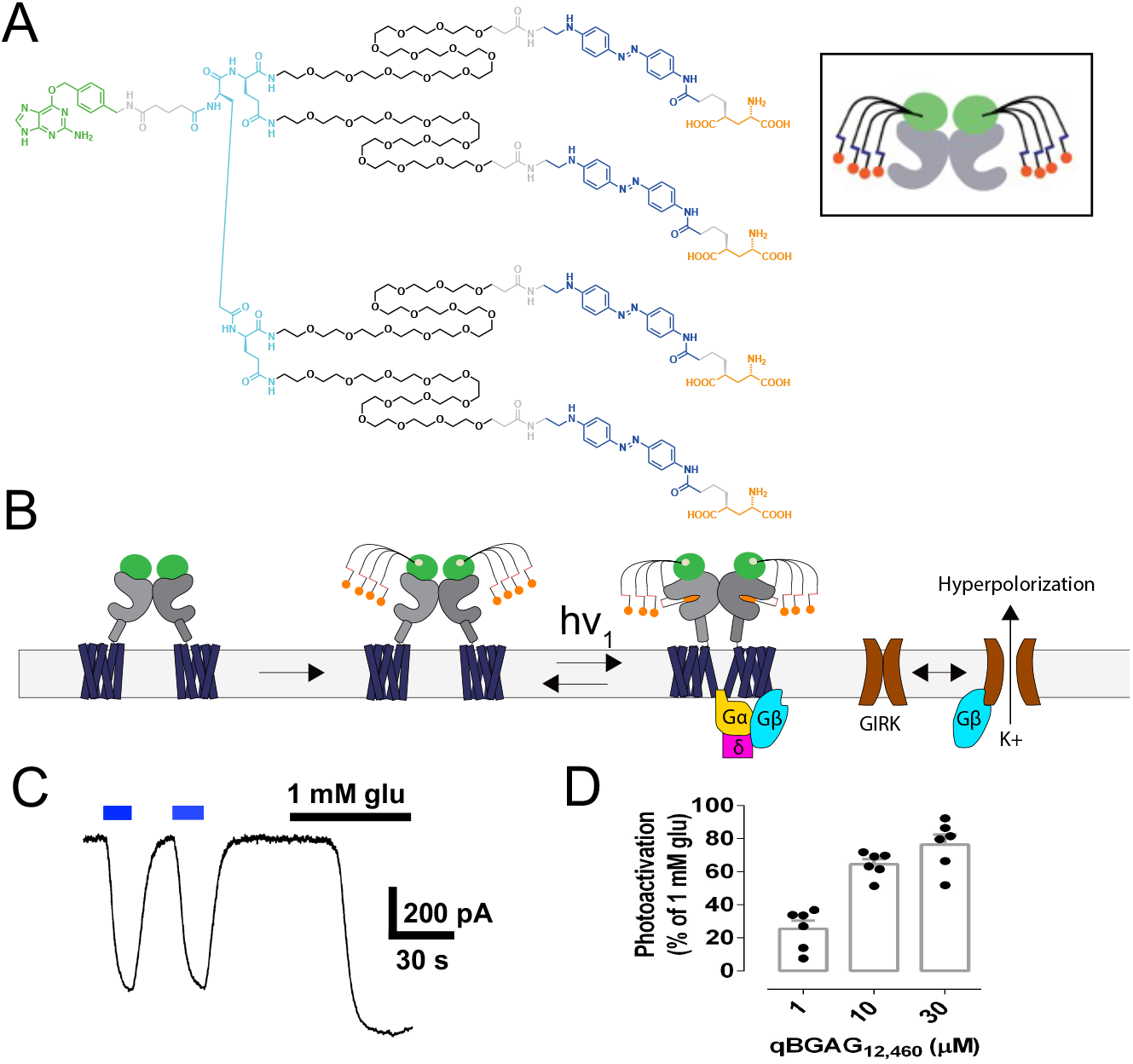
Four-branched ^4x^BGAG_12,460_ photoactivates SNAP-mGluR2. **(A)** Chemical structure of ^4x^BGAG_12,460_ with Benzylguanine SNAP-attachment moiety (green), PEG-linkers (black), azobenzene (blue) and L-glutamate ligand (orange) color labeled. Inset cartoon depicts attachment of one ^4x^BGAG_12,460_ to the SNAP fused to the mGluR2 N-terminal in the mGluR2 dimer. **(B)** Schematic of SNAP-tagged mGluR2 without (left) and with (middle) covalently tethered ^4x^BGAG_12,460_ photoswitches. Photoactivation of mGluR2 (right) leads to binding of G_i_/Gβγ and liberation of Gβγ, which binds to and activates and opens the GIRK channel (brown) leading to K+ efflux (and hyperpolarization). **(C)** Whole-cell patch-clamp trace in HEK293T cell shows inward current through GIRK channels (120 mM [K^+^]_ext_, V_H_ = 80mV) illustrates that SNAP-mGluR2 is activated by the *cis*-configuration of ^4x^BGAG_12,460_ under blue (470 nm) light and relaxes spontaneously back to the *trans*-configuration in the dark., leading to receptor and channel deactivation. Photoactivation of ^4x^BGAG_12,460_ –SNAP mGluR2 is reproducible and efficient (~80% max) when compared with application of saturating (1 mM) glutamate. **(D)** Summary bar graph of ^4x^BGAG_12,460_-SNAP mGluR2 photoactivation efficacy in HEK293T cells labeled for 30 min at different concentrations of ^4x^BGAG_12,460_ (n=6 cells per concentration) expressed as a percentage of activation by saturating (1 mM). glutamate.

^4x^BGAG_12,460_ was applied to HEK293T cells, which co-expressed SNAP-mGluR2 and the G protein activated inward rectifier potassium channel (GIRK), at 1 - 30 μM for 1 hr. Photoactivation of the G_i_-coupled mGluR2 was measured through the Gβγ activation of the G protein activated inward rectifier K^+^ (GIRK) channel, which generates a K^+^ current that we observed as an inward current in high external K^+^ at a negative holding potential in whole cell patch clamp recording (**Figure 1B, C**). Maximal optical activation was obtained with labeling at 10 and 30 μM, yielding a photo-current that was ~75% the size of the current activated by saturating (1 mM) glutamate (**Figure 1C, D-source data 1**). This level of photo-activation is ~2x higher than achieved with the single branched BGAG_12,460_^10^, confirming a recent report ^20^, and suggesting that ^4x^BGAG_12,460_ could increase the light sensitivity of restored vision.

To test the function of ^4x^BGAG_12,460_ in the retina, we turned to the same vector construct used in our earlier study^10^, with SNAP-mGluR2 under the control of the promoter of the human Synapsin 1 gene, which is preferentially expressed in retinal ganglion cells (RGCs) of mice and humans^23,24^. The plasmid, which included inverted terminal repeats (ITRs) and the woodchuck hepatitis virus posttranscriptional regulatory element (WPRE) (Methods), was packaged into the AAV2(4YF) capsid. Delivery into the eye of ≥3-month-old *rd1* mice was *via* intravitreal injection. As shown earlier,^10^, AAV2(4YF):hSyn-SNAP-mGluR2 drives strong expression of SNAP-mGluR2 in the RGC layer of the *rd1* retina, as visualized by labeling of SNAP with a membrane-impermeable BG-dye instead of with the BGAG photoswitch.

Six to ten weeks after intravitreal AAV injection, we injected 2 μl of 1 mM of ^4x^BGAG_12,460_ in PBS into the ~5 μl volume mouse eye, to produce a concentration of ~290 μM in the vitreous. To assess the light evoked response, SNAP-mGluR2 expressing retinas were removed from *rd1* mice and mounted on a multi-electrode array (MEA) (**Figure 2A, B**). Due to photoreceptor degeneration, the retinas of untreated *rd1* adult mice (>12wks), and ones expressing SNAP-mGluR2 in RGCs, do not have recordable light-evoked responses at the intensities that we tested ^10^. However, *rd1* mouse retinas expressing SNAP-mGluR2 in RGCs, from animals that had been injected intravitreally with ^4x^BGAG_12,460_ in PBS 2-4 days earlier, showed light-evoked suppression of firing, followed by a transient excitatory rebound at the cessation of illumination (**Figure 2C-source data 2**), a response that resembles the photoreceptor cell driven response of OFF-RGCs in the *wt* retina ^25,26^. With application of progressively shorter light pulses, the amplitude of the rebound excitation declined more than did the inhibition elicited during light exposure, leaving substantial inhibition even in response to a light pulse of only 25 ms (**Figure 2D-source data 2**). Notably, despite the 40-fold reduction in the number of photons delivered as flash duration was reduced from 1000 to 25 ms, the inhibitory response maintained the same fast kinetics. This suggests that ^4x^BGAG_12,460_:SNAP-mGluR2 may support motion vision even in dim light.

**Figure 2:**
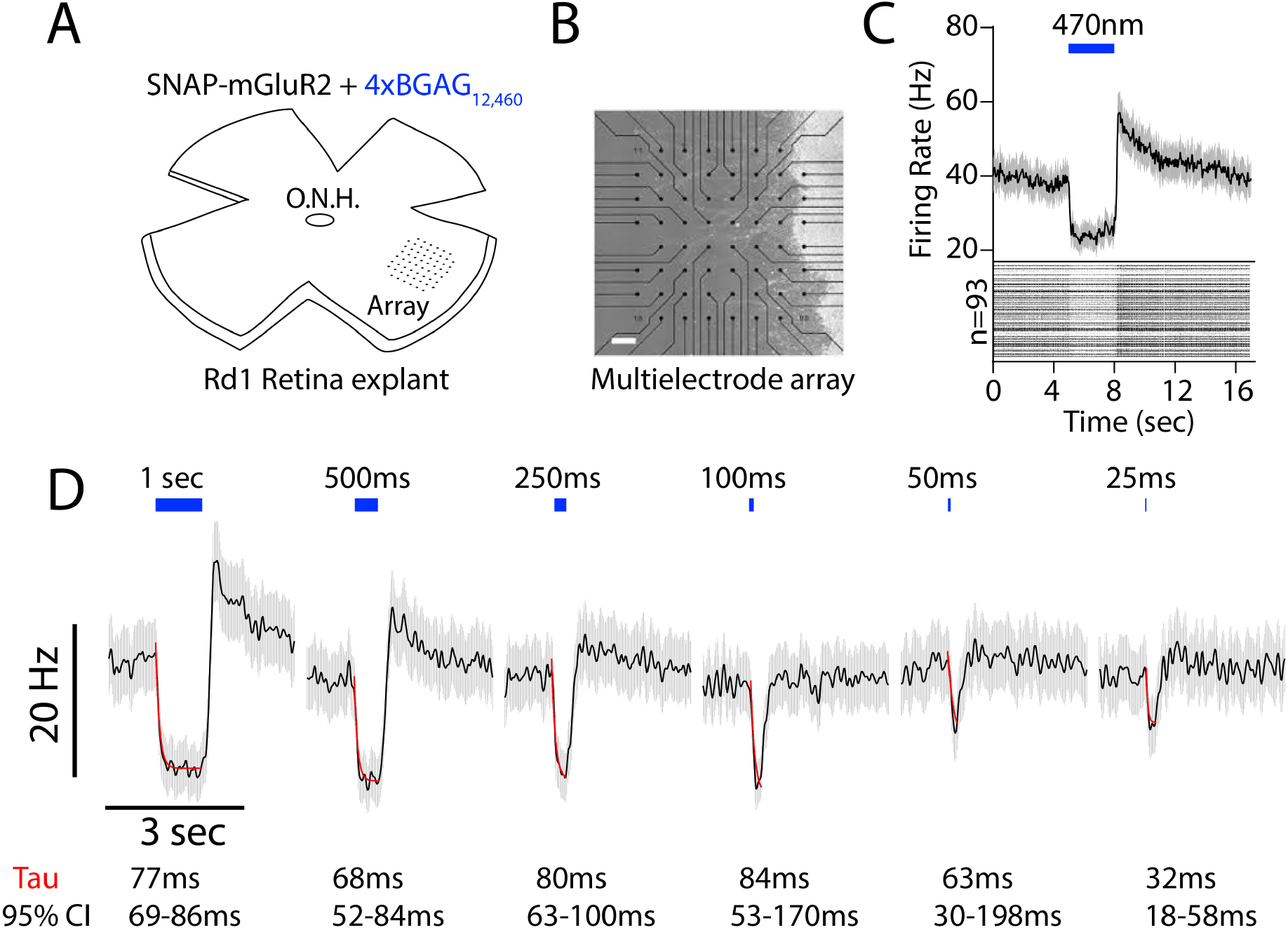
^4x^BGAG_12,460_ restores fast retinal light response to the *rd1* mouse retina. (A, B) Illustration of *rd1* mouse retina expressing SNAP-mGluR2 in RGCs, labeled wth ^4x^BGAG_12,46,_ excised and mounted on a 60-channel transparent multielectrode array (MEA). **(C)** Photoactivation of ^4x^BGAG_12,460_-SNAP-mGluR2 in RGCs triggers a fast suppression of spontaneous firing followed by a rebound excitation when the light is turned off. (Bottom) Raster plot of response in 93 RGCs in *rd1* retina with ^4x^BGAG_12,460_-SNAP-mGluR2 to a 3 sec flash of illumination (blue = 472nm) (each cell shows average response to 5 pulses of light. (Top) Average response of the *93* RGCs shown in the raster (bottom). **(D)** Dependence of ^4x^BGAG_12,460_:SNAP-mGluR2 RGC light response on flash duration (*n*=93). Population average firing shows detectable responses down to 25 ms illumination duration. Single exponential fit (red) superimposed on inhibitory component of responses shown. Time constants from fits with 95% confidence intervals (95% CI) indicated below. **(C, D)** *n* = sorted cells. Mean (black) ±SEM (gray).

### Branched BGAG restores high-sensitivity light aversion

Wildtype (C57) mice with intact visual function naturally avoid illuminated spaces, however mice with retinal degeneration, such as *rd1*, do not change their behavior in response to differences in illumination and cannot perform this light/dark discrimination^10,16,19,27^. To assess the ability of *rd1* animals expressing SNAP-mGluR2, which has been derivatized with BGAG, to distinguish light from dark, we employed a 2-chamber shuttle box, with an open doorway between the chambers (**Figure 3A**) and an iPad-mini installed on the far wall of each chamber, one of which was set to black and the other to white. The illuminated LCD display was set to one of four room light intensities, ranging from dim to standard lighting: 0.2, 5, 25 or 88 μW/cm^2^ (*i*.*e*. 10^γ^ photons cm^−2^ s^−1^, where γ = 11.7, 13.1, 13.8 or 14.3, respectively). In this apparatus, normally sighted mice avoid the illuminated chamber, whereas vision impaired *rd1* mice spend equal amounts of time in the two chambers ^15^. On days 5-7 after injection 2 μl of 1 mM ^4x^BGAG_12,460_ in PBS, mice spent more time in the dark chamber than in the illuminated chamber when the illumination was at 5, 25 or 88 μW/cm^2^ but not when it was 0.2 μW/cm^2^ (**Figure 3B-source data 3**).

**Figure 3.**
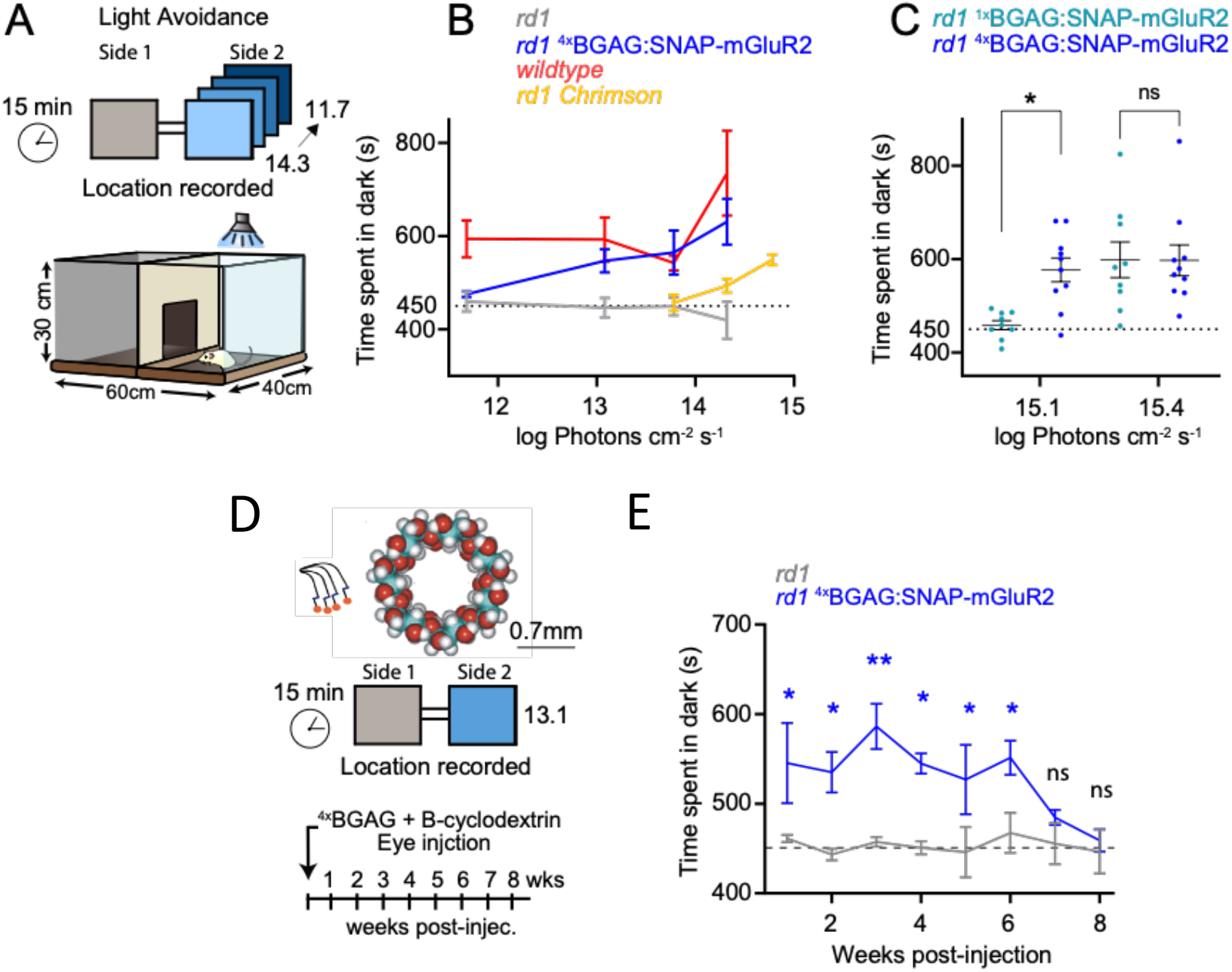
^4x^BGAG_12,460_ in β-CD restores high sensitivity light avoidance for weeks. **(A)** Schematic of light/dark box for assessing light avoidance using a consumer LCD tablet that is turned off in one chamber (grey) and illuminated at range of light intensities in other chamber (shades of blue). **(B)** Amount of time spent in the dark compartment when the illuminated chamber is lit with 10^γ^ photons cm^−2^ s^−1^ of blue light (445 nm) (where γ = 11.7, 13.1, 13.8, 14.3 or 14.78 log photons cm^− 2^ s^−1^; equivalent to 0.2, 5, 25, 88, and 250 uW cm-^2^) for four animal groups: *rd1* control (gray; *n* = 7, 8, 9, 11, and 6 mice, respectively), *rd1* mice with expressing SNAP-mGluR2 in RGCs and injected with ^4x^BGAG_12,460_ in β-CD (blue; *n*= 6, 8, 8, 6 mice, respectively), *rd1* mice with expressing ChrimsonR in RGCs (orange; *n*= 6,13,10 mice, respectively), and wildtype (C57) mice (red; *n* = 6, 5, 4, 5 mice, respectively). **(C)** Amount of time spent in the dark compartment when the illuminated chamber is lit with 10 ^γ^ photons cm^−2^ s^−1^ of blue light (445 nm) (where γ = 15.1 or 15.4 photons cm^−2^ s^−1^ equivalent to 0.5 and 1.0 mW cm-^2^) for *rd1* mice with expressing SNAP-mGluR2 in RGCs and injected with either unbranched (single ligand) BGAG_12,460_ (teal; *n* = 9, 9 mice, respectively) or ^4x^BGAG_12,460_ (blue; *n*= 10, 10 mice, respectively) in β-CD. Statistical significance assessed using Student’s unpaired *t*-test; ***: p=<0.001. **(D)** (Top) Illustration of a beta cyclodextrin (β-CD) (right) that is combined with ^4x^BGAG_12,460_ (left) as a method of slow-release photoswitch delivery to the eye. (Below) schematic of light avoidance behavioral stimuli repeated over an 8-week period following a single injection to yield a vitreal concentration of ~290 μM ^4x^BGAG_12,460_ combined with ~1mM β-CD. **(E)** Light avoidance of *rd1* mice (gray; n= 6 mice) compared to *rd1* mice expressing SNAP-mGluR2 (blue; n=8 mice) over an 8-week period following a single intravitreal injection of ^4x^BGAG_12,460_ in β-CD. Light avoidance performed at 88 μW cm^−2^ (14.3 log photons cm^−2^ s^−^). Mean±SEM. Statistical significance assessed using Student’s unpaired *t*-test: *: p=<0.05, **: p=<0.01.

To contextualize this sensitivity, we compared this performance to *rd1* mice expressing ChrimsonR, an enhanced channelrhodopsin^28^ that is currently in phase 1/2a clinical studies in patients. We found that a white screen at an intensity of 88 μW cm^−2^, the maximal brightness of the iPad-mini, evoked weak photo-aversion (**Figure 3B-source data 3**). We therefore turned to a brighter Tripltek display and found that at its maximal brightness of 250 μW cm^−2^ (14.8 log photons cm^−2^ s^−1^), photo-aversion in *rd1* mice expressing ChrimsonR in RGCs was similar to that obtained at 25 μW/cm^2^ (13.8 log photons cm^−2^ s^−1^) with ^4x^BGAG_12,460_: SNAP-mGluR2 in RGCs (**Figure 3B-source data 3**), representing an ~10-fold higher sensitivity of ^4x^BGAG_12,460_: SNAP-mGluR2 in RGCs. To determine the effect on visual sensitivity of increasing BGAG branch number, and hence photo-ligand number, we compared light avoidance behavior in *rd1* mice expressing the same SNAP-mGluR2 in RGCs but injected with ^1x^BGAG_12,460_ that places only one photoswitchable agonist per binding site. *Rd1* mice expressing SNAP-mGluR2 in RGCs, injected with either ^1x^BGAG_12,460_ or ^4x^BGAG_12,460_ photoswitches exhibited strong light avoidance at 1mW cm^−2^ (15.4 log photons cm^−2^ s^−1^) (**Figure 3C-source data 3**). However, at a light intensity of 0.5mW cm^−2^ (15.1 log photons cm^−2^ s^−1^) there was no difference between *rd1* animals expressing SNAP-mGluR2 in RGCs that were injected with ^1x^BGAG_12,460_ and *rd1* control mice, indicating absence of avoidance to this level of light. These results suggests that the 4-branched branched BGAG provides an ~100-fold increase in sensitivity over single-branched BGAG.

We wondered if formulating ^4x^BGAG_12,460_ in β-cyclodextrin (**Figure 3D**), an excipient commonly used for topical ocular delivery, whose variants are used for various routes of delivery of diverse drugs to many tissues,^29^ would extend the restoration of light perception, as we observed earlier with intravitreal delivery of single armed BGAG_12,460_ mixed with cyclodextrin ^10^. Two to four days after intravitreal injection of 2 μl of 1 mM ^4x^BGAG_12,460_ in 0.5% β-cyclodextrin (for an intra-ocular concentration of ~290 μM ^4x^BGAG_12,460_ in ~1mM β-cyclodextrin; See Methods) we observed restored photo-aversion in *rd1* animals expressing SNAP-mGluR2, which lasted for 6 weeks after a single intravitreal injection (**Figure 3E-source data 3**). After this period, treated animals returned to spending equal time in the illuminated and dark chambers, similar to untreated *rd1* animals, providing a functional time course of restored photosensitivity *in vivo*.

Thus, ^4x^BGAG_12,460_ activates SNAP-mGluR2 strongly enough to support detection of moderate indoor light, with light aversion occurring at ~1/3^rd^ of the maximum brightness of a standard computer display. Moreover, formulation in β-cyclodextrin extends this action for weeks after a single injection, as observed earlier with unbranched (monovalent) BGAG_12,460_ ^10^, suggesting that this formulation could represent a general method for extending the action of BGAG-like PORTLs (Photoswitchable Orthogonal Remotely Tethered Ligands).

### ^4x^BGAG_12,460_:SNAP-mGluR2 restores novel object exploration

Having observed that ^4x^BGAG_12,460_:SNAP-mGluR2 enables light discrimination with dim displays, we wondered how it would operate for the recognition of three-dimensional objects under incidental light at room light intensity. To address this, we employed an open field arena that is commonly used to test novel object recognition and exploratory behavior ^30,31^. Sighted mice naturally avoid open spaces and maintain proximity to walls of their environment. Exploratory excursions from these places of safety can be motivated by novel objects^19^. Although mice employ multiple sensory modalities during exploration, vision is critical for spatial navigation^32^. The square arena contained two distinct novel objects that were thoroughly cleaned to remove olfactory cues. The mouse was placed against the arena wall, at a sufficient distance from the objects, which themselves were sufficiently separated, to minimize accidental encounters. We video recorded untreated *rd1* mice and *rd1* mice that expressed SNAP-mGluR2 RGCs, which had been injected 5 weeks earlier with ^4x^BGAG_12,460_ in β-cyclodextrin, as above. Mouse movement was tracked for 10 minutes the first time that they were placed into the arena (**Figure 4A**). We measured the latency to exploration of the novel objects and the distance and velocity that the animals traveled. Untreated *rd1* mice had long latencies to reach the objects, whereas *rd1* mice expressing SNAP-mGluR2, which had been injected with ^4x^BGAG_12,460_ in β-cyclodextrin, had significantly shorter latencies to the exploration of the objects (**Figure 4B, C-source data 4**). Untreated *rd1* mice locomoted more slowly and covered shorter distances within the observation time than did *rd1* mice with SNAP-mGluR2 and spent a larger fraction of time along the wall of the arena (**Figure 4D, E-source data 4**). These results suggest that ^4x^BGAG_12,460_:SNAP-mGluR2 provides visually impaired mice with a retinal degeneration with the ability to detect objects visually under incidental light at the intensity of room light.

**Figure 4:**
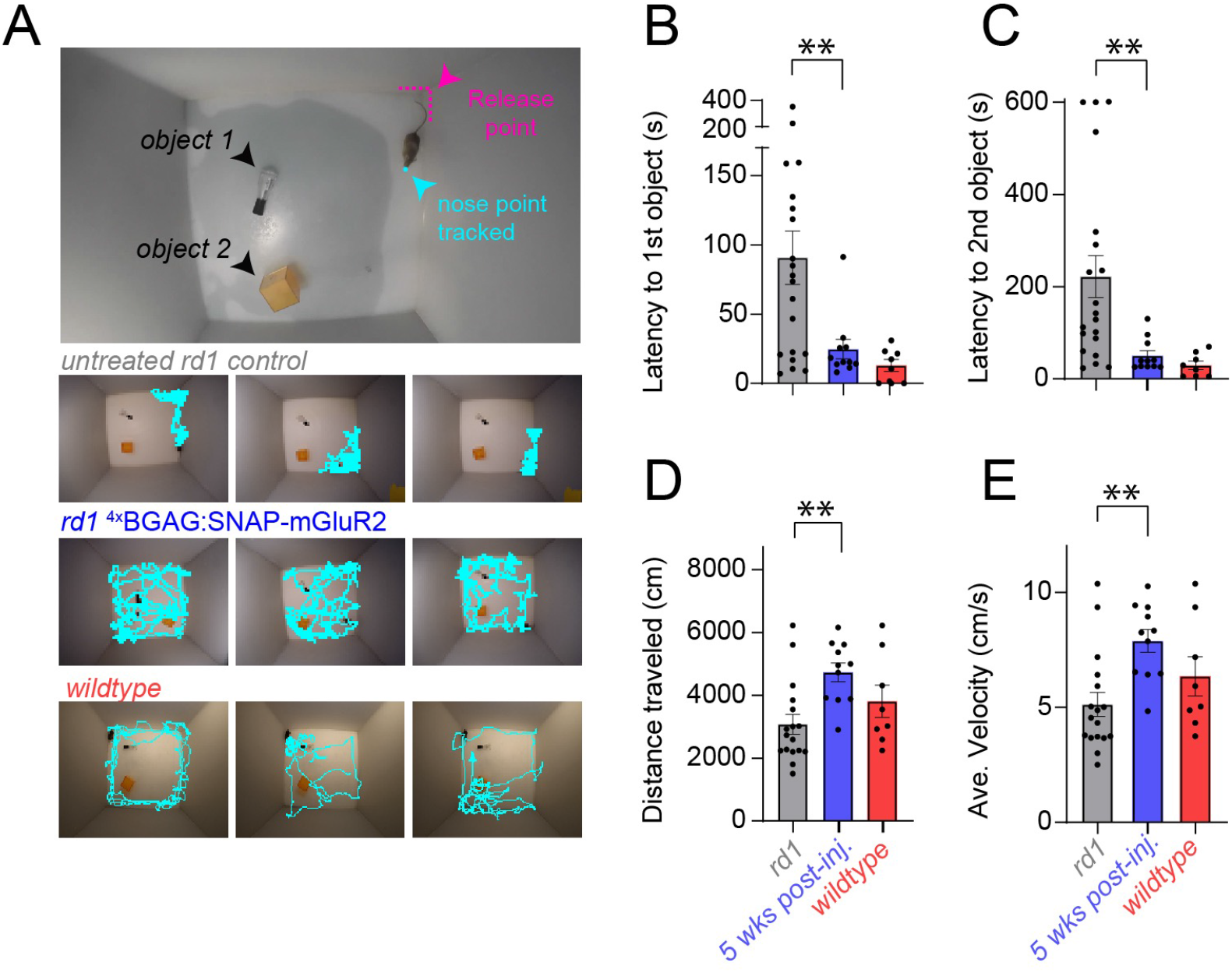
Novel object recognition restored in indoor light 5 weeks after injection of ^4x^BGAG_12,460_ in β-CD. **(A)** Open field behavioral arena containing two novel objects with traces of the first minute locomotion track from 3 representative animals per condition: untreated r*d1* mice (top), ^4x^BGAG_12,460_:SNAP-mGluR2 expressing *rd1* mice (middle), and wildtype (C57) mice (bottom). (B-E) Bar graphs displaying components of novel object exploration in *rd1* mice (gray; n=17), *rd1* mice expressing SNAP-mGluR2, 5 weeks after injection of ^4x^BGAG_12,460_ in β-CD (blue; n=11), and wildtype mice (red; n=8). Mean±SEM Statistical significance assessed using Student’s unpaired *t*-tests: **: p=<0.01 **(B, C)** Latency to explore the objects. Latency to first object (B) and cumulative latency to the second object (C). **(D, E)** Total distance traveled (D) and average velocity (E) of mice during 10 min of exploration.

### ^4x^BGAG_12,460_:SNAP-mGluR2 restores high-sensitivity and high-acuity line pattern recognition

We next asked whether ^4x^BGAG_12,460_:SNAP-mGluR2 would support patterned vision. We tested the ability of treated and untreated *rd1* mice to discriminate between two different line patterns in a learned negative association task. The task is carried out in a 2-chamber arena, with a conductive floor in each chamber which can apply an aversive mild foot shock (**Figure 5A**). Each chamber had an iPad-mini at its far end, which displayed a unique line pattern, consisting of black-on-white parallel vertical lines at one of two spacings. The mouse was habituated to the chamber for one day and trained for two days, during which a shock was given in one of the chambers. The mouse was then tested on a final day (recall phase) when no foot shock was given and only the displays were shown (**Figure 5A, top**). The room was kept dark to avoid room reference visual cues and the aversive and non-aversive cues were switched on the day of recall to ensure that any location bias would work against the preference for the non-aversive display.

**Figure 5:**
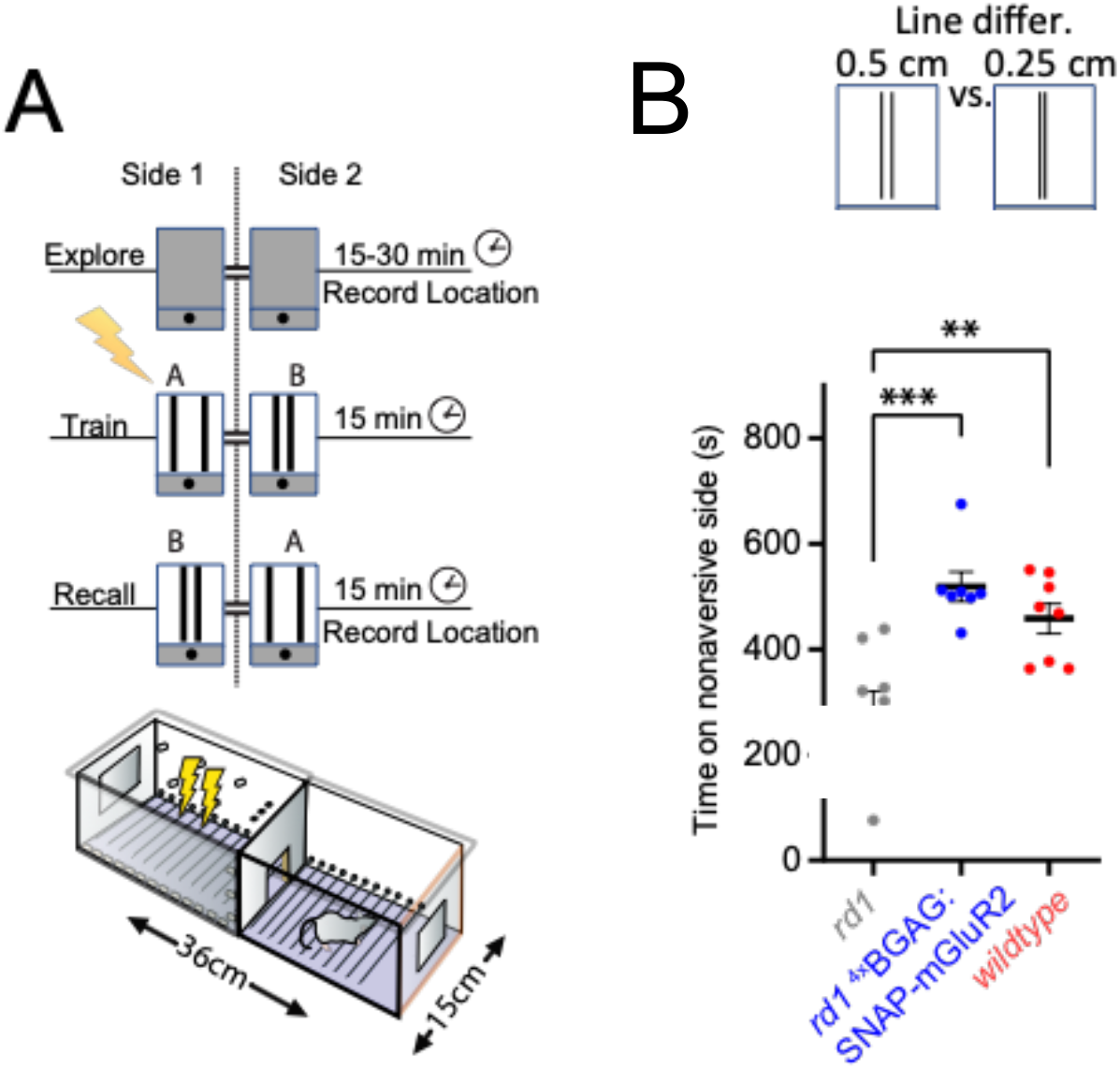
^4x^BGAG_12,460_:SNAP-mGluR2 in RGCs restores learned pattern discrimination to *rd1* mice. **(A)** (Top) Schematic of displays used in pattern discrimination aversion association task. (Bottom) 2-chamber behavioral arena connected by doorway, with mounted LCD tablets on far walls and metal grill floors for applying mild foot shock. Day 1: mice habituated to chamber. Days 2-3: shock in one chamber during display of distinct pattern in each chamber. Day 4: Recall tested (as time spent on non-aversive side) in absence of shock with light patterns reversed from training period to avoid location bias. **(B)** (Top) Schematic of displays used in high acuity pattern discrimination aversion association task. High acuity learned pattern discrimination between vertical lines spaced 0.25 cm *versus* 0.5 cm apart. Untreated *rd1* mice (gray; n=8), *rd1* mice with ^4x^BGAG_12,460_-SNAP-mGluR2 (blue; n=7). Wildtype (C57) mice (red; n=8). Mean ± SEM. Statistical significance assessed using one-way ANOVA test with Holm-Sidak correction for multiple comparisons; **: p=<0.01, ***: p=<0.001.

Provided they have sufficient visual acuity to distinguish between the two-line patterns, sighted animals avoid the side with the line pattern that had been previously associated with the foot shock, whereas untreated *rd1* mice display a location bias by favoring the side associated with the aversive cue during recall; as described previously.^10,19^ The LCD screens in the two chambers each displayed a pair of identical vertical lines, 0.5 cm wide. In one chamber the separation between the lines was 0.25 cm and in the other chamber it was 0.5 cm. Given the distance from the displays to the doorway separating the chambers (18 cm), which we treated as the point of decision, these line separations distances are equivalent to 0.75° and 1.5° of arc, respectively (see Methods), around the limit of acuity that has been estimated for wildtype mouse vision ^33,34^. We found that *rd1* mice with ^4x^BGAG_12,460_:SNAP-mGluR2 in RGCs were able to distinguish between these displays and spend more time in the chamber containing the display that had not been associated with the foot shock during training (**Figure 5B-source data 5**). These observations suggest that ^4x^BGAG_12,460_:SNAP-mGluR2 in RGCs restores vision with an acuity that is similar to normal mouse vision.

### ^4x^BGAG:SNAP-mGluR2 restores superior line pattern recognition in motion

The kinetics of the restored light response are expected to affect the ability to track a visual pattern and recognize it in motion. Whereas the rate of rise of current generated by light-activation of channelrhodopsin depends on the intensity of the light, so that dim illumination generates slower responses, we found ^4x^BGAG_12,460_-labeled SNAP-mGluR2 in RGCs to have relatively constant inhibitory light response kinetics across light doses (**Figure 2D, Tau inhibition**). We therefor predicted that ^4x^BGAG_12,460_-labeled SNAP-mGluR2 would provide *rd1* mice with the ability to resolve patterns in motion. To address this hypothesis, we measured the ability of *rd1* mice to perform pattern recognition with moving lines. For comparison, we tested *rd1* mice expressing ChrimsonR ^28^.

As ^4x^BGAG_12,460_-labeled SNAP-mGluR2 endows higher photosensitivity than ChrimsonR (**Figure 3B**), we compared performance of rd1 mice with ^4x^BGAG_12,460_- labeled SNAP-mGluR2 in RGCs using the iPad-mini display at 88 μW cm^−2^ (14.3 log photons cm^−2^ s^−1^) to rd1 mice with ChrimsonR in RGCs using the Tripltek display at 250 μW cm^−2^ (14.8 log photons cm^−2^ s^−1^). *Rd1* mice expressing ChrimsonR in RGCs were able to distinguish between line pairs separated by 1 *versus* 6 cm when the display was immobile (0°/s) at a level that was similar to that of *rd1* mice with ^4x^BGAG_12,460_:SNAP-mGluR2 in RGCs, as well as that of *wt* mice (**Figure 6B-source data 6**). When the line pairs moved horizontally across the screen during both the training and testing periods (**Fig. 6A**), *rd1* mice with ^4x^BGAG_12,460_:SNAP-mGluR2 in RGCs performed as well as *wt* mice at up to 12 cm/s (36°/s) (**Figure 6B-source data 6**). In contrast, *rd1* mice expressing ChrimsonR in RGCs performed at the level of the untreated *rd1* mice (*i*.*e*. were unable to distinguish between the line patterns) at either 12 cm/s (36°/s) or 4 cm/s (12°/s), the slowest speed tested (**Figure 6B-source data 6**). These observations indicates that ^4x^BGAG_12,460_:SNAP-mGluR2 supports pattern recognition in motion at moderate room light intensity, while ChrimsonR even at optimal outdoor light levels, does not allow pattern recognition in motion.

**Figure 6:**
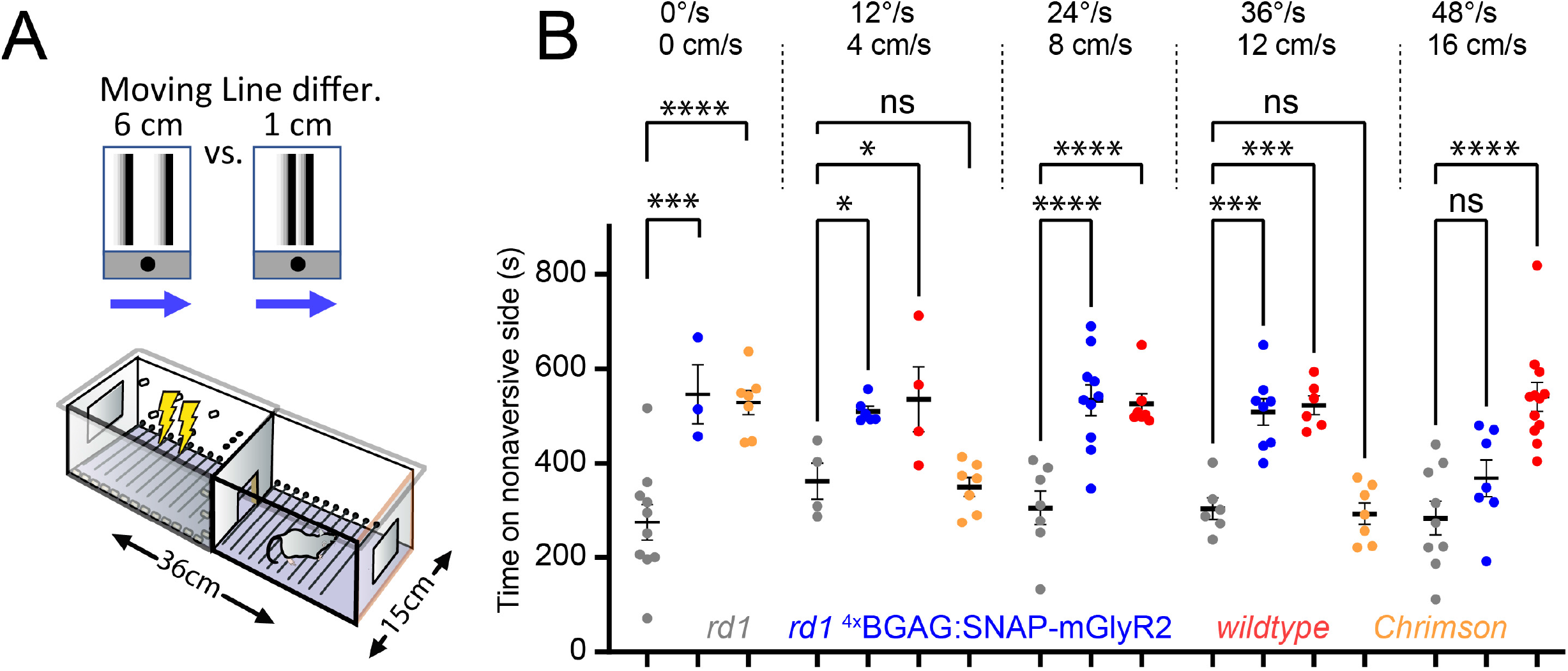
^4x^BGAG_12,460_:SNAP-mGluR2 restores moving pattern discrimination. **(A)** Schematic of discrimination between pairs of lines separated by 6 versus 1 cm while the lines move to the right at varying speed in the aversion association task. **(B)** Pairs of vertical lines, 2 cm in width, move left to right at speeds of 0 (immobile), 4, 8, 12 or 16 cm s^−1^ (equivalent to 0, 12, 24, 36 or 48°/s). Time spent in chamber with display that was not associated during training with foot shock is significantly longer at all but fastest speed of motion in *rd1* mice with ^4x^BGAG_12,460_:SNAP-mGluR2 (blue; n=3, 6, 10, 8, 7, respectively) compared to untreated *rd1* mice (gray; n=10, 4, 7, 6, 9, respectively). *Rd1* mice expressing ChrimsonR in RGCs also prefer chamber with display that was not associated during with foot shock, but only when the display is immobile (0°/s; orange; n = 7), and performed like untreated *rd1* mice when the lines moved at 12 or 36°/s (orange; n=7, 7, respectively). Wildtype mice (red) shown for comparison (n=4, 7, 6, 12 at 12, 24, 36 or 48°/s, respectively). Mean±SEM. Statistical significance assessed using one-way ANOVA test; *: p=<0.05, ***: p=<0.001, ****: p=<0.0001.

## Discussion

Most inherited retinal degenerations cause progressive death of rod and cone photoreceptor cells eventually leading to complete blindness. However, downstream retinal neurons are largely spared, providing a potential target for restorative photosensitivity through the installation of light-activated signaling protein. Several years ago, we demonstrated vision restoration to the *rd1* mouse model using intravitreal AAV to introduce into RGCs a gene encoding a SNAP-tagged version of the Gi-coupled glutamate-activated GPCR mGluR2 (SNAP-mGluR), followed by the intravitreal injection of the agonist BGAG_12,460_, which attaches covalently to SNAP and activates the receptor in response to light ^10^. This photoswitch attaches covalently to SNAP and activates the receptor in response to light. The restored visual function required bright stimuli and so was limited in function to the intensity of outdoor light intensities. In the current study, we boosted the sensitivity to light, so that the restored vision could operate at indoor light intensities. We find that the system restores object recognition, acuity at the level of normal vision and line pattern recognition even in rapid motion.

### Supra-linear increase in light sensitivity with branched BGAG

To increase sensitivity, we employed a newly designed BGAG photoswitch where the SNAP attachment moiety (BG) is coupled to multiple PEG branches, each bearing its own photoswitchable azobenzene-glutamate (AG). Once attached to SNAP, the branched BGAG places as many photo-agonists as there are branches next to each glutamate binding pocket, increasing the chance that light will photoisomerize one of the AGs into the configuration that can bind and activate the receptor. We used a scaffold with four branches, ^4x^BGAG_12,460_, which increases by ~2-fold the efficacy of SNAP-mGluR2 activation, as measured by photocurrent from the co-expressed G protein activated inward rectifier GIRK channel in HEK293 cells^20^. Here we test ^4x^BGAG_12,460_ for the first time in vision restoration. Strikingly, when SNAP-mGluR2 is expressed in the RGCs of the *rd1* mouse retina, the sensitivity to light was increased by ~40-fold with ^4x^BGAG_12,460_ compared to the single branched BGAG_12,460_, much more than the ~2-fold increase in efficacy seen in HEK293 cells. This large increase visual sensitivity may arise downstream of receptor activation. Our earlier pharmacological analysis suggested that the mGluR2 effector channel in the RGCs of the *rd1* mouse is not the GIRK channel, but KCNQ4 ^10^. KCNQ4 is a member of the Kv7 family of voltage-gated potassium channels, which, like the GIRK channel, is activated by the Gβγ that is released by Gi-coupled receptors, such as mGluR2 ^35–38^. Whereas the GIRK channel begins to open when a single Gβγ binds to one of its four subunits, and progressively increases opening as more Gβγs bind to additional subunits^39^ Kv channels typically must have all 4 subunits activated to open^40^. This could account for the supra-linearity in the relationship between mGluR2 activation and the hyperpolarizing potassium current in RGCs. With multi-branched BGAG on SNAP-mGluR2 in RGCs, the *rd1* mouse was able to perform novel object recognition under incidental room light and to differentiate between patterns of lines shown on a standard LCD computer display.

### High acuity of restored pattern discrimination

We used an aversive association task to assess the acuity of the restored vision and found that *rd1* mice with ^4x^BGAG_12,460_:SNAP-mGluR2 in RGCs could distinguish between parallel lines that are separated by 0.75° *versus* 1.5° of arc. This result is remarkable as this acuity approaches the reported limit of *wt* vision in mouse^41^. This performance stands out compared to the acuity levels of vision restoration using enhanced channelrhodopsin-2 variants, which fall short of *wt* performance^12^. However, studies of acuity mediated by channelrhodopsins have used reflexive head tracking in response to moving gradients in an optokinetic drum, distinct from learned discrimination with an immobile line pattern, preventing direct comparison. Future analysis should employ a common approach, while accounting for differences in sensitivity.

### Fast signaling kinetics in RGCs and line pattern recognition in motion

A key to restoring vision in motion is for the optogenetic system to have sufficiently fast to allow for a rapid retinal refresh. While photo-stimulation of channelrhodopsin can evoke currents that rise to peak in a few milliseconds in response a bright flash of light (typically ~1 mW/mm^2^) the channels gate proportionately more slowly when illumination is at lower intensity^42^. In contrast, with either ^4x^BGAG_12,460_:SNAP-mGluR2 in the RGCs of the *rd1* mouse, flashes of dim (room light intensity) light elicit fast responses, and reducing the number of photons by a 40-fold reduction in light flash duration (from 1000 ms to 25 ms) does not slow the photo-current kinetics (Fig. 2D). This suggests that the branched BGAG:SNAP-mGluR2 will provide better motion vision at indoor light levels than channelrhodopsins.

Indeed, we find that *rd1* mice with ^4x^BGAG_12,460_:SNAP-mGluR2 in RGCs can distinguish between pairs of lines separated by different distances, even when the lines move at 48°/s and perform as well as *wt* mice at 36°/s. In contrast, ChrimsonR, one of the enhanced channelrhodopsins^28,42^, and the first optogenetic system demonstrated to restore vision in a patient^11^, did not work in motion, even at one third of the displacement speed. ChrimsonR activation accelerates as light intensity increases^42^. In our experiments, ChrimsonR was tested with displays that were ~3-fold brighter than those used for ^4x^BGAG_12,460_:SNAP-mGluR2, but apparently insufficient in brightness. However, under bright light, or with intensifying goggles, ChrimsonR may well signal rapidly enough to support vision in motion.

Our findings demonstrate that ^4x^BGAG_12,460_:SNAP-mGluR2 provides superior motion vision over one of the more promising microbial channelrhodopsins. This advance is promising for practical vision under natural circumstances, such as navigating the environment, examining objects, reading and using a computer. Consistent with this, ^4x^BGAG_12,460_:SNAP-mGluR2 supported detection and exploration of objects under incidental illumination at the intensity of room light.

### Summary of 2-component chemical optogenetics for vision restoration

A key requirement of 2-component chemical optogenetics vision restoration is the need to follow the intravitreal viral delivery of the gene encoding SNAP-mGluR2 with intravitreal delivery of the synthetic ^4x^BGAG_12,460_ photoswitch. Free ^4x^BGAG_12,460_ is cleared from the eye and the ^4x^BGAG_12,460_-conjugated receptors turn over. To sustain vision restoration, ^4x^BGAG_12,460_ injection would need to be repeated at regular intervals. We find that adding β-cyclodextrin to the of ^4x^BGAG_12,460_ formulation extends the restoration of vision to 6 weeks after injection, as shown earlier for the unbranched BGAG_12,460_^10^. This frequency of injection is similar to that used with intravitreal injection of anti-VEGF antibodies (*Lucentis* or Eylea) to treat neovascular macular degeneration and may be possible to further extend by other slow-release delivery methods. The added burden of repeated photoswitch injection is offset by the advantages of high sensitivity and high acuity and function in motion. The use of mGluR2, which is normally expressed in the retina, may reduce the risk of immune reaction from introduction of a foreign gene product in the eye. Moreover, because the chemical photoswitch washes out, it could allow for use of improved photoswitches, as they become available, as well as for adjustment of dose to optimize the therapy to the needs of individual patients. Finally, retinal diseases differ in progression and, for patients with partial vision, it may be preferable to select a therapeutic whose action is temporary, rather than a completely genetic optogenetic therapy that, once introduced, cannot be turned off.

## Methods

### Animals, AAVs and photoswitches

Mouse experiments were performed under the express approval of the University of California Animal Care and Use Committee. Wildtype mice (C57BL/6 J) and rd1 mice (C3H) were purchased from the Jackson Laboratory and bred in house. Animals were housed on a 12-h light/dark cycle with food and water ad libitum. cDNA encoding SNAP-mGluR2 was inserted in an established viral backbone under control of the human Synapsin promoter (hSyn-1) and packaged in the AAV2/2-4YF capsid^10^. The plasmid included flanking inverted terminal repeats (ITRs), a polyA tail and the woodchuck hepatitis virus posttranscriptional regulatory element (WPRE) to yield: AAV2(4YF):ITR-hSyn-SNAP-mGluR2-polyA-WPRE-ITR. The vector was sequenced to validate the integrity of the construct. AAV at a concentration of 10^11^–10^12^ viral genomes was injected in a 2 μl volume to the vitreous of the *rd1* mouse eye via microinjection to both eyes. rAAV injections were at p30–p60 and *in vivo* and *in vitro* experiments were conducted at p90– p160. AAVs were produced as previously described.^10,19^

4xBGAG_12,460_ was synthesized as described previously.^19^ For injections in PBS, stock solution of 100 mM ^4x^BGAG_12,460_ (L-diastereomer) in 100% pharmaceutical grade DMSO (Cryoserv; Bioniche Pharma) was diluted 1:100 in sterile PBS for a final working solution of 1 mM in 1.0% DMSO. This PBS stock was delivered in a 2 μl volume to the ~5 μl volume vitreous of the rd1 mouse eye via microinjection, for a final vitreal concentration of ~290 μM ^4x^BGAG_12,460_. For formulation in β-cyclodextrin, the 100 mM ^4x^BGAG_12,460_ stock in 100% DMSO was diluted 1:25 in sterile PBS to 4 mM in 4% DMSO and then further diluted 1:4 in 16 mM β-cyclodextrin for a final stock concentration of 1 mM ^4x^BGAG_12,460_ in 4 mM β-cyclodextrin. This β-cyclodextrin stock was delivered in a 2 μl volume to the ~5 μl volume vitreous of the rd1 mouse eye via microinjection, for a final vitreal concentration of ~290 μM ^4x^BGAG_12,460_ and ~1.14 mM β-cyclodextrin.

### Electrophysiology

To assess the efficacy of ^4x^BGAG_12,460_:SNAP-mGluR2 photoactivation, we measured currents induced in GIRK channels. HEK293T cells were seeded sparsely onto 18 mm coverslips and maintained in DMEM (Invitrogen) with 10% fetal bovine serum on poly-L-lysine-coated coverslips at 37°C and 5% CO_2_. The cells were transiently transfected overnight with Lipofectamine 2000 with the following DNA constructs: SNAP fused to the N-terminus of rat mGluR2 (SNAP-mGluR2; 0.7 μg), a homotetramerizing mutant of GIRK1 (F137S; 0.7 μg), and tdTomato (0.1 μg). All constructs were cloned into mammalian expression vectors. Sixteen to 24 hours after transfection, the HEK293T cells were incubated with 1, 10 or 30 μM ^4x^BGAG_12,460_ for 60 minutes in the dark at 37°C in standard extracellular buffers. The cells were then washed and voltage clamped in whole-cell configuration in an extracellular solution that contained 120 mM KCl, 25 mM NaCl, 10 mM HEPES, 2 mM CaCl_2_, and 1 mM MgCl_2_, pH 7.4. Glass pipettes with a resistance of 3-7 MΩ were filled with intracellular solution containing 120 mM gluconic acid δ-lactone, 15 mM CsCl, 10 mM BAPTA, 10 mM HEPES, 1 mM CaCl_2_, 3 MgCl_2_, 3 mM MgATP, pH 7.2. Cells were voltage clamped to −80 mV using an Axopatch 200A (Molecular Devices) amplifier. Glutamate was applied using a gravity-driven perfusion system and illumination with blue light (445 nm; ~4 mW/mm^2^) was applied to the entire field of view using a Lambda DG4 (Sutter) through a 20x objective. pClamp software was used for both data acquisition and control of illumination. The selection criteria for electrophysiological experiments were that a cell (i) expresses the fluorescent protein transfection marker, and (ii) responds to glutamate, indicating the presence of mGluR2 and GIRK. We did not exclude individual cells unless a recording was of poor quality (e.g., unstable baseline). Multi-electrode array recordings were performed on excised retina using a perforated 1060 system (Multi Channel Systems). Spikes were exported from voltage traces using MC_Rack, sorted in Plexon Offline sorter and analyzed and graphed in Neuroexplorer (Plexon) and MATLB (MathWorks), as previously described.^10,19^

### Behavior

#### Passive avoidance open field test

For the passive avoidance open field test a two-compartment (light and dark) shuttle box (Colbourn Instruments) arranged to allow the mouse to move freely through a small opening that connects the two compartments. For wt (C57), untreated rd1, rd1 expressing SNAP-mGluR2 in RGCs and labeled with ^4x^BGAG_12,460_, and rd1 expressing ChrimsonR in RGCs, the light compartment was illuminated by a consumer iPad-mini generation 4 centered over the light compartment and display intensity was either 0.2, 5, 25 or 88 μW/cm^2^ at the display screen. For rd1 expressing ChrimsonR in RGCs, one brighter illumination was used with 250 μW/cm^2^ at the display screen (Tripltek model T82) centered over the light compartment. Mice were placed in the light compartment and were given a maximum of 3min to discover that there is a second compartment. A 15-min trial began when they crossed into the dark compartment, and time spent in the light was recorded. This previously established approach has been described ^10^.

#### Learned active avoidance task

For the active avoidance task, a two-compartment, shuttle box (Colbourn Instruments) arranged to allow the mouse to move freely through a small opening that connects the two compartments, however, unlike the passive avoidance open field task, the floor in each chamber is conductive and used to apply an aversive mild foot shock (Fig. 6A). Each chamber had an LCD display at its far end, which displayed a unique line pattern. For rd1 mice with ^4x^BGAG_12,460_:SNAP-mGluR2 in RGCs, the displays were iPad-minis at maximum brightness of 88 μW/cm^2^ at the display screen. For rd1 mice with ChrimsonR in RGCs, the displays were Tripltek model T82s at maximum brightness of 250 μW/cm^2^ at the display screen. The stimuli consisted of black-on-white parallel vertical lines, 0.5 cm in width. The lines were spaced at different distances, reported in cm on the display screen as well as degrees of arc, as calculated based on the 18 cm distance between each screen and the point of decision at the doorway between the chambers. Our experiments compared spacing of 0.25 vs 0.5 cm (0.75° vs 1.5°; Fig. 5B) or 1 vs 6 cm (3° vs 18°) in either a still display or with the lines moving at four different speeds = 4, 8, 12 or 16 cm s^−1^ (12°, 24 °, 36 ° or 48 ° s^−1^, respectively; Fig. 6) (see figure legends for details). Mice were trained for two days, during which a shock was given in one of the chambers. On the third day (recall phase), no foot shock was given, and only the displays were shown. The room was kept dark to avoid room reference visual cues and the aversive and non-aversive cues were switched on the day of recall to avoid location bias. This previously established approach was described in detail earlier ^19^.

#### Object exploration task

For the object exploration task, two objects were placed in a 50 × 50 cm open field box. Animals were positioned in the empty box and allowed to explore freely over the course of 10 min. The following day, two novel objects (object 1 & 2) were placed in the box and animals were again positioned along the wall of the box and allowed to explore freely for 10 min while the arena was filmed continuously. Unbiased analysis was performed using Noldus Technology Ethosvision XT v13.5. Videos were analyzed for distance traveled (cm), the velocity of travel (cm/s), and the latency (s) to arrive at the first and second object. This previously established approach was described earlier ^19^. These experiments were done under light from above with an intensity of 170 μW/cm^2^at the floor of the chamber.

#### Calculation of acuity

We calculated the acuity of restored vision using paired line discrimination. iPad-minis, placed on the far wall of each chamber of the 2-chamber shuttle box, displayed pairs of black vertical lines on a white background at the maximal intensity of the display of 88 μW/cm^2^. The line width and length were the same in the two displays, but the spacing between the lines differed. In Figure 5, the immobile displays showed 0.5 cm wide lines separated by either 0.25 cm or 0.5 cm. In Figure 6, a larger spatial difference was presented in a moving display, with 1 cm wide lines separated by either 1 cm or 6 cm that moved at 4-16 cm s^−1^. We calculated the angle of arc based on the distance of 19 cm from the decision point, which we defined as the doorway between the chambers, to the display. Perimeter = 2Πr = 119 cm. 360°/119cm = 3°/cm. 0.25 cm = 0.75°; 0.5 cm = 1.5°.

## Author Contributions

A.H., M.B., P.D. and E.Y.I. designed the experiments, A.H., M.B., and P.D. performed the experiments and analyzed the results, with input from E.Y.I. Synthesis of ^4x^BGAG_12,460_ was by J.B., viral synthesis and cloning by M.V., HEK cell recordings by P.D., retinal MEA recordings by M.B. and behavioral experiments by A.H. and J.L. The paper was written by E.Y.I. with input from all of the authors.

## Acknowledgments

We thank Cherise Stanley, Roya Hosseini, and Erik Lasker for technical assistance. The work was supported by the National Institutes of Health (R01EY028240; E.Y.I. and J.G.F.), the Foundation Fighting Blindness, USA (TA-GT-0818-0745-UCB; J.G.F.) and Vedere Bio II, Inc. (E.Y.I. and J.G.F.).

## Data Availability

All data generated during this study are included in the manuscript and supporting source data files.

## Materials Availability

All molecular and chemical reagents will be made available upon request.

